# Mannan molecular sub-structures control nanoscale glucan exposure in *Candida*

**DOI:** 10.1101/215632

**Authors:** Matthew S. Graus, Michael J. Wester, Douglas W. Lowman, David L. Williams, Michael D. Kruppa, Jesse M. Young, Harry C. Pappas, Keith A. Lidke, Aaron K. Neumann

**Affiliations:** Department of Pathology, University of New Mexico, Albuquerque, NM 87131; Department of Mathematics and Statistics, University of New Mexico, Albuquerque, NM 87131; Center of Excellence in Inflammation, Infectious Disease and Immunity, Quillen College of Medicine, East Tennessee State University, Johnson City, TN 37684; AppRidge International, LLC, Telford, TN 37690; Department of Surgery, Quillen College of Medicine, East Tennessee State University, Johnson City, TN 37684; Department of Biomedical Sciences, Quillen College of Medicine, East Tennessee State University, Johnson City, TN 37684; Department of Physics and Astronomy, University of New Mexico, Albuquerque, NM 87131

## Abstract

N-linked mannans (N-mannans) in the cell wall of *Candida albicans* are thought to mask β-(1,3)-glucan from recognition by Dectin-1, contributing to innate immune evasion. Lateral cell wall exposures of glucan on *Candida albicans* are predominantly single receptor-ligand interaction sites and are restricted to nanoscale geometries. *Candida* species exhibit a range of basal glucan exposures and their mannans also vary in size and complexity at the molecular level. We used super resolution fluorescence imaging and a series of protein mannosylation mutants in *C. albicans* and *C. glabrata* to investigate the role of specific N-mannan features in regulating the nanoscale geometry of glucan exposure. Decreasing acid labile mannan abundance and α-(1,6)-mannan backbone length correlated most strongly with increased density and nanoscopic size of glucan exposures in *C. albicans* and *C. glabrata*, respectively. Additionally, a *C. albicans* clinical isolate with high glucan exposure produced similarly perturbed N-mannan structures and exhibited similar changes to nanoscopic glucan exposure geometry. We conclude that acid labile N-mannan controls glucan exposure geometry at the nanoscale. Furthermore, variations in glucan nanoexposure characteristics are clinically relevant and are likely to impact the nature of the pathogenic surface presented to innate immunocytes at dimensions relevant to receptor engagement, aggregation and signaling.

## Introduction

The cell wall of *Candida* species yeasts is an essential feature that provides structural support and protection. There are three major polysaccharide components: mannan, β-(1,3;1,6)-glucan and chitin(Chattaway et al., 1968). These components are organized into two layers, an inner layer primarily composed of chitin and β-glucan, and an outer layer mostly composed of cell wall proteins decorated with N- and O-linked glycosylations known as mannans (Klis et al., 2001)(Poulain et al., 1978)(Ruiz-Herrera et al., 2006)(Nguyen et al., 1998)(Netea et al., 2008). Previous evidence linking elevated glucan exposure in *C. albicans* mutants with impaired cell wall biosynthesis has implicated N-mannan in restricting immunogenic β-glucan exposure at the cell wall surface, presumably via a steric masking effect (Wheeler et al., 2008).

N-mannans consist of an N-glycan core, an α-(1,6)-mannoside backbone and side chains that contain α-(1,2) or (1,3) linked oligomannosides. Serotype A *C. albicans* may also possess side chain β-(1,2) or (1,3) linked oligomannosides. Additionally, acid labile β-mannoside moieties of variable length may be appended to N-mannan side chains via phosphodiester linkages (Cutler JE, 2001)(West et al., 2013)(Hall et al., 2013)(Hall and Gow, 2013). β-(1,2) or (1,3) linked oligomannosides have also been identified in *C. albicans* as part of the phospholipomannan moiety of the cell wall (Trinel et al., 2002). The exact structure of N-mannan varies between *Candida* species and is dependent on the expression of a complex network of mannan biosynthesis, trafficking and cell wall remodeling genes (Shibata et al., 2012).

The outer layer of the cell wall, the point of contact between the yeast and the immune system, is predominantly composed of N-mannans with punctate exposures of β-glucan (Gantner et al., 2005). These components act as pathogen-associated molecular patterns (PAMPs), which are recognized by the immune system through pattern recognition receptors (PRRs) (Gow and Hube, 2012)(Medzhitov and Janeway, 1997)(Gow et al., 2012). C-type lectins (CtLs) are a class of PRRs that include DC-SIGN, mannose receptor (CD206) and Mincle which bind to N-mannan, TLR4 which binds O-linked mannans, and Dectin-1 which binds β-glucans (Gow et al., 2012)(Netea et al., 2006)(Gow et al., 2007)(Bugarcic et al., 2008)(Wells et al., 2008).

Clustering is a prominent feature of PRRs and a common mechanism of regulating receptor activity (Inoue and Shinohara, 2014). Previously, we reported changes in CtL nanocluster geometry during fungal particle recognition and the presence of nanoscale ligand patterns on *C. albicans* cell walls (Itano et al., 2012)(Lin et al., 2016). We have also reported that *C. albicans* glucan is sparsely accessible on lateral yeast and hyphal cell walls, consisting of single glucan/Dectin-1 interaction sites as well as larger (~40 nm diameter) exposures (Lin et al., 2016). Recent studies on reactive oxygen species (ROS) generation from glucan-coated particles of varying size (50, 200 or 500 nm diameter) (Goodridge et al., 2011) suggest that ligand presentation geometry may influence Dectin-1 clustering and signaling. Multimerization of Dectin-1 upon ligand engagement is thought to be important for signal transduction via the receptor’s hemITAM domains. Therefore, the nanoscale geometry of glucan exposure is likely to impact Dectin-1 signaling and generation of innate immune anti-fungal responses.

In this report, we sought to better define nanometric glucan exposure geometries by determining structural features of *C. albicans* and *C. glabrata* N-mannans that regulate glucan exposure geometry at the molecular level.

## Results

### Phagocytic response and β-glucan exposure varies between species

We began the study by measuring phagocytosis efficiencies between *C. albicans* (SC5314) and *C. glabrata* (ATCC2001) on dendritic cells (DCs). DCs co-cultured with ATCC2001 had a higher phagocytic efficiency when compared to the DCs co-cultured with SC5314 (Fig. 1A).

**Figure 1.**
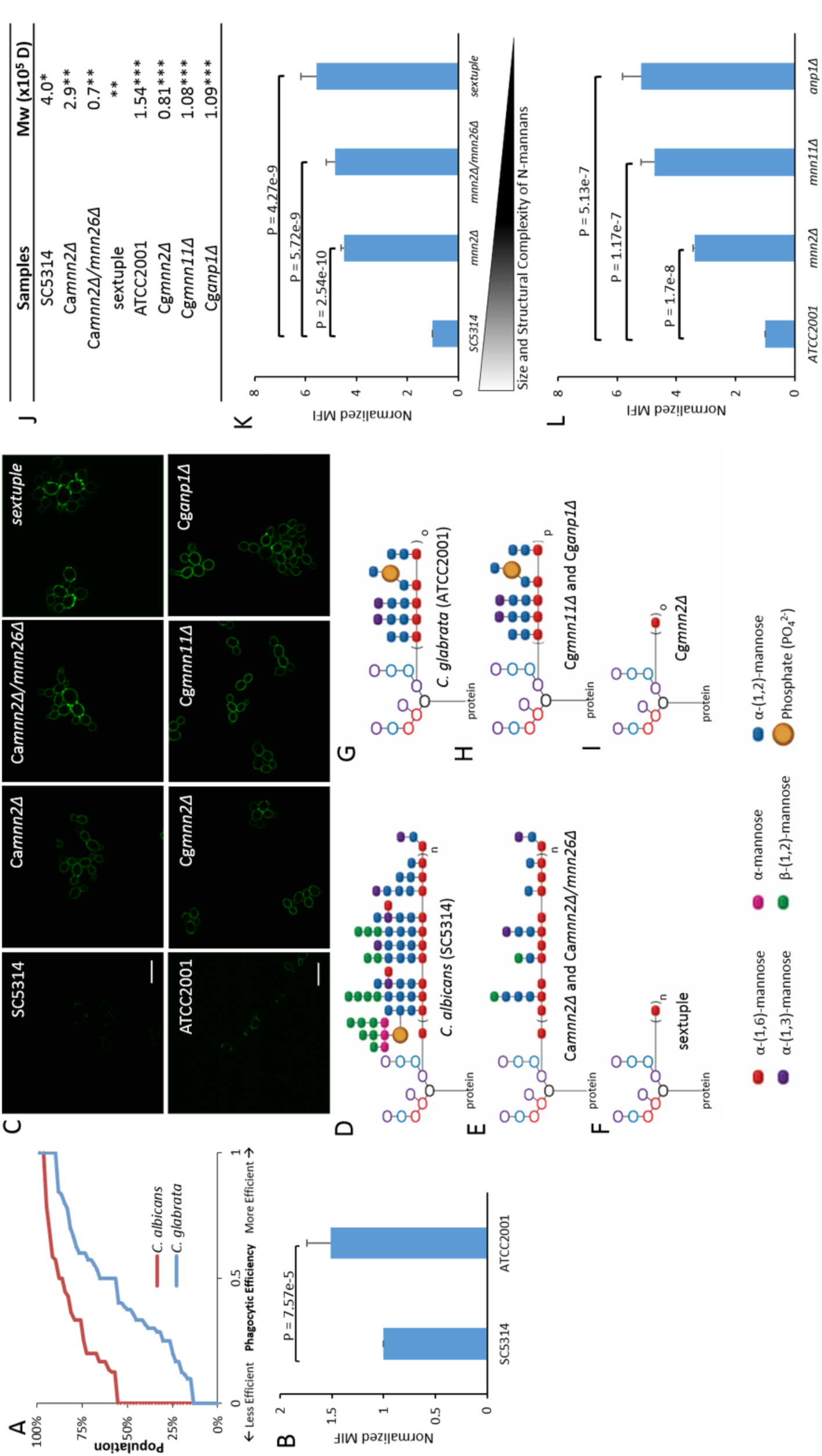
N-mannan structure correlates with the amount of β-glucan exposed. A) Phagocytic efficiency of the distribution for populations of primary human DCs incubated with either SC5314 or ATCC2001. DCs' counted n ≥ 50. B,K,L) Comparison of the integrated intensity of β-glucan measured from flow cytometry on yeasts. Statistical significance was determined by a single-tailed t-test with n ≥ 20000 gated single cell per strain. Values and error bars displayed in B,K,L are presented as normalized medians and standard deviations. C) Representative images of yeast strains with varying amounts of β-glucan exposed. Scale bar = 10 µm. D-I) Schematic presentation of N-mannan structures for either type strains or mannosylation mutants. Subscripts n, o, and p correspond to an unspecified number of repeats > 1, and o > p. E and F structures have been adapted from Hall et al. 2013. G-I structures have been adapted from West et al. 2013. J) Molecular weights of type strains and mannosylation mutants. * Data from Kruppa, et al, 2011, ** Data from Hall et al. 2013 sextuple data was not available, *** Data from West et al. 2013.

Glucan exposure has been linked to phagocytosis in other studies(Hardison and Brown, 2012),(Graus et al., 2014), suggesting that differences in the amount of glucan exposed could relate to the changes in phagocytic efficiencies. Although the *C. albicans* cell wall contains 10% more β-glucan and 17% less mannose than *C. glabrata*(de Groot et al., 2008) by dry mass, flow cytometry determined that β-glucan exposure on *C. glabrata* was 1.5-fold greater than on *C. albicans* (Fig. 1B,C). This suggests that glucan exposure and DC phagocytic uptake are inversely correlated with cell wall glucan abundance and led to the hypothesis that the molecular structure of mannan may be the major determinant of glucan exposure. Consistent with this hypothesis, previous studies reported the molecular weight of *C. albicans* N-mannans to be 2.6-fold greater than that of *C. glabrate* (West et al., 2013)(Hall et al., 2013). In addition, the molecular structure of *C. glabrata* mannan is lower in structural complexity relative to *C. albicans* (Shibata et al., 2012).

### Comparison of *C. albicans* and *C. glabrata* strains' N-mannan structure

Gene deletion mutants that have significant and known defects in mannan biosynthesis were used to explore the possibility that size and molecular complexity of N-mannans determines the ability of those mannans to sterically hinder access of receptors to glucan, thus regulating surface glucan exposure. We began with the CaMNN2 gene family mannosylation mutants previously characterized in Hall *et al.* 2013 (Hall et al., 2013). This gene family is responsible for normal abundance and complexity of N-mannan side chains (Hall et al., 2013)(Rayner and Munro, 1998), and deletion mutant strains of the CaMNN2 family exhibit altered mannan structure. The *C. albicans* parental strain was compared to the mannosylation mutants *mnn2*Δ, *mnn2*Δ*/mnn26*Δ, and a sextuple mutant (all CaMNN2 gene family deleted). In general, as the number of CaMNN2 genes deleted increased, the molecular weight and structural complexity of the mannans declined, but because of the higher multiplicity of MNN2 family members in *C. albicans*, a greater variety of mannan structures are produced relative to *C. glabrata*, wherein only one MNN2 gene has been characterized. Specifically, Ca*mnn2*Δ and Ca*mnn2*Δ*/mnn26*Δ were previously found to be structurally similar by NMR with regard to the types of side chain structures present and lacking. CaMMN2 family mutant strains exhibited marked reduction in acid labile mannan fraction and reduced abundance and complexity of side chains relative to the parental strain, along with diminishing Mw as the severity of mutation increased (Fig. 1D,E,J). The *C. albicans* sextuple MNN2 family deletion mutant produces the most severely perturbed mannan, containing only a unsubstituted α-(1,6)-mannose backbone. This makes the sextuple mutant’s mannan most structurally analogous to the Cg*mnn2∆* mannan investigated below (Fig. 1F,I). There is no direct analog between Ca*mnn2*Δ and Ca*mnn2*Δ*/mnn26*Δ mutant’s mannans and the *C. glabrata* mannans investigated below as a result of their more extensive side chain defects and more extreme diminution of molecular weight. Based on previous work(Hall et al., 2013), we hypothesized that the negatively charged acid labile mannan as well as the large complex side chains are important N-mannan features for masking β-glucan.

### Analysis of glucan exposure in *C. albicans* strains

Measurement by flow cytometry with the *C. albicans* mannosylation mutants showed that total glucan exposure increases as N-mannan size and structural complexity decreases, resulting in a 4.5 to 5.5-fold increase in total exposures relative to SC5314 (Fig. 1C,K).

Processes that unmask glucan change not only the total amount of glucan exposed, but also the nanostructural features of glucan presentation to innate immunoreceptors like Dectin-1(Lin et al., 2016). direct Stochastic Optical Reconstruction Microscopy (dSTORM) was used to explore the unmasking process. dSTORM data was quantified with the Hierarchical Single Emitter Hypothesis Test (H-SET) and two different clustering algorithms to measure the geometry and characteristics of the exposures. H-SET is able to correct common artifacts in dSTORM imaging arising from multiple localizations of individual probes, allowing accurate structural representation of nanoscale objects (Lin et al., 2016). The clustering algorithms hierarchical-based clustering and Voronoï tessellation-based segmentation (Levet et al., 2015) define clusters and measure nanoscopic cluster characteristics.

dSTORM imaging revealed that the majority of exposures in all strains are single probe binding events with the sextuple mannosylation mutant having the highest density of single probe events (Fig. 2A-D; SuppFig. 1A-C). When compared to SC5314, all mannosylation mutants possess significantly more glucan multi-exposures that are larger in diameter (Fig. 2E,F; SuppFig. 1D,E). In the most severely affected sextuple mutant, glucan exposure density increased by 4.5-fold (relative to SC5314) and median exposure size increased from 14 nm (SC5314) to 21 nm (sextuple mutant). We conclude that N-mannan side chain feature deficiencies affect glucan unmasking by increasing both the number of exposures on the cell wall and the sizes of the exposures. The side chain characteristics altered in these mutant strains appear to control glucan exposure nanostructure through diminished density of side chains and the loss of acid labile mannan.

**Figure 2.**
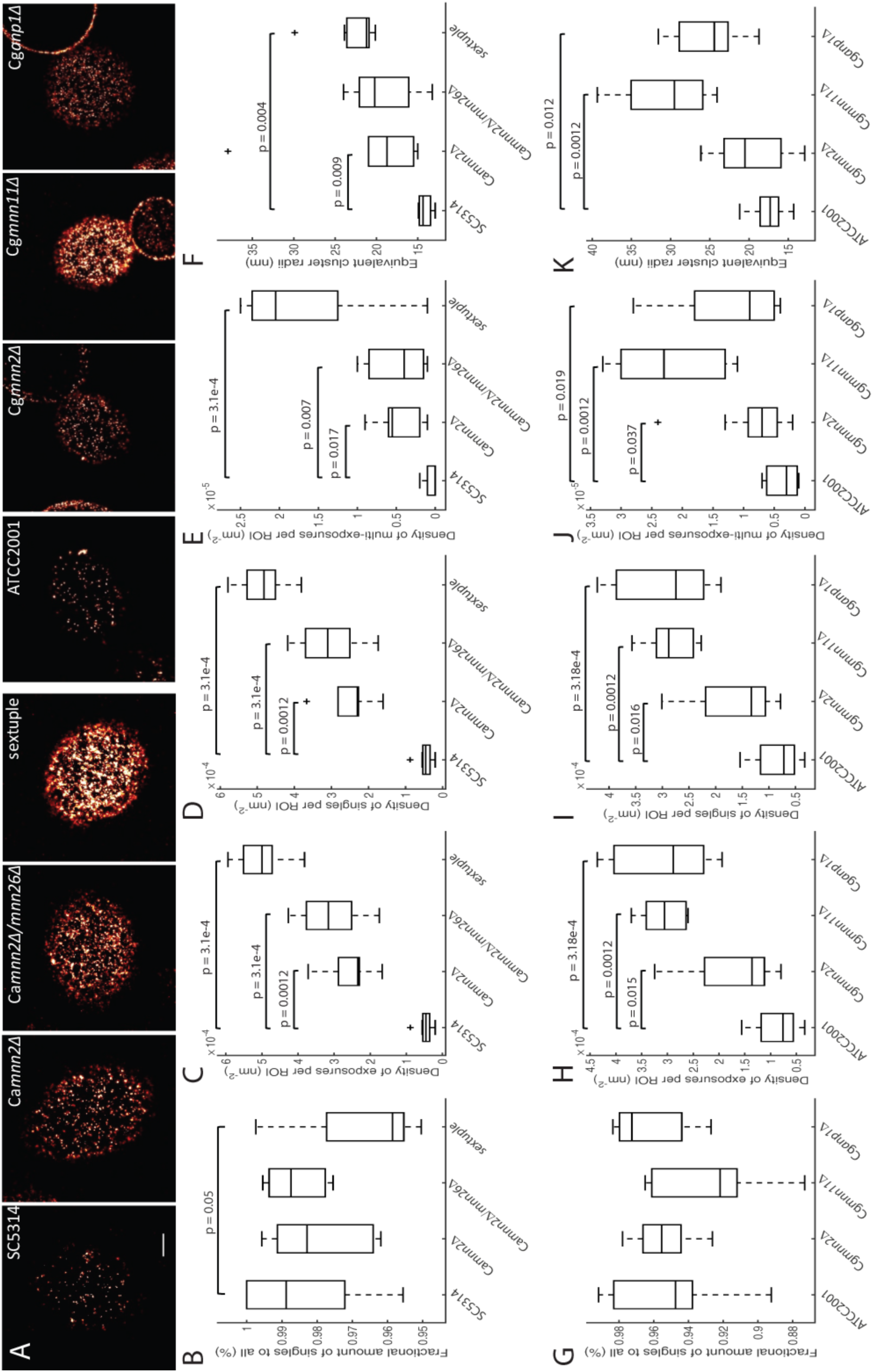
Nanoscopic β-glucan exposure characteristics are determined by N-mannan structure. A) Representative dSTORM images of type strains and mannosylation mutants. Scale bar = 750 nm. B-F) Hierarchical clustering quantification of nanoscopic glucan exposure geometries on C. albicans strains. G-K) Hierarchical clustering quantification of nanoscopic glucan exposure geometries on C. glabrata strains. Box plots represent population of events found in ROI. Statistical significance was determined by Mann-Whitney U test with n ≥ 6 for each strain.

### Analysis of glucan exposure in *C. glabrata* strains

To study how N-mannan structure affects glucan exposure in the relatively small mannans of *C. glabrata* mannosylation mutants from West *et al.* 2013(West et al., 2013) were utilized. The authors demonstrated alterations in mannan structure of Cg*anp1∆*, Cg*mnn11∆* and Cg*mnn2∆* relative to the parental strain, ATCC2001. N-mannans in the mannosylation mutants are smaller than that of the parental strain, with Cg*mnn2∆* mannan having the largest decrease in molecular weight(West et al., 2013) (Fig. 1J). Both Cg*anp1∆* and Cg*mnn11∆* strains have a reduced length of the mannan α-(1,6)-mannose backbone, but retain the α-(1,2) and α-(1,3)-mannose side chains in reduced abundance because of the shorter backbone (Fig. 1H). Cg*mnn2∆* mannan has an unsubstituted α-(1,6)-mannose backbone similar to the sextuple mutant (Fig 1F,I). Furthermore, Cg*mnn11∆* and Cg*anp1∆* retain the acid labile fraction of its mannan, but this negatively charged constituent is missing in Cg*mnn2∆*. We hypothesized Cg*mnn2*Δ would have the largest amount of β-glucan exposed due to the lack of space-filling side chains that are likely be important for steric masking of underlying glucan.

All three mannosylation mutants exhibit increased amounts of glucan exposed relative to the parental *C. glabrata* strain. Surprisingly, Cg*mnn2∆* exhibited the smallest increase in glucan exposure, while Cg*mnn11∆* and Cg*anp1∆* had much larger increases in comparison to ATCC2001 with fold increases that ranged from 3.3 to 5 (Fig. 1C,L). The reduction of the α-(1,6)-mannose backbone length in Cg*mnn11∆* and Cg*anp1∆* unexpectedly plays a more significant role on N-mannan’s ability to mask glucan efficiently. To understand how the shortening of the α-(1,6)-mannose backbone affects unmasking, dSTORM imaging was used to examine the *C. glabrata* mannosylation mutants.

Similar to the *C. albicans* mannosylation mutants, the majority of exposures identified in the *C. glabrata* data set are single probe binding events (Fig. 2A,G-I; SuppFig. 1F-H). Interestingly, shortening of the α-(1,6)-mannose backbone has the greatest effect on increasing the density of nanoscale glucan exposures, with a 1.5-fold increase in clustered glucan exposures for Cg*mnn2∆* relative to ATCC2001, and that the median exposure size increases from 21 nm (Cg*mnn2∆*) to 29 nm (Cg*mnn11∆*). The data suggests that the ability to extend a side chain-decorated mannan backbone in space is the most critical parameter for overall glucan masking and the density of glucan nanoexposures in *C. glabrata*.

### Analysis of glucan exposure in *Candida* clinical isolate strains

We extended the study by investigating glucan exposure in recent clinical isolates of *C. albicans* and *C. glabrata*. Most *C. albicans* and *C. glabrata* clinical isolates have low levels of glucan exposure, similar to the SC5314 and ATCC2001 type strains (SuppFig. 2A and data not shown). However, one *C. albicans* isolate (“TRL035”) and one *C. glabrata* isolate (“TRL031”) were found that exhibited a total glucan exposure 3.1-fold greater than SC5314 and 1.5-fold greater than ATCC2001, respectively (SuppFig. 2A-D).

Due to its unusual cell wall phenotype, TRL035’s identity was confirmed as *C. albicans* by several methods. In sum, classification of this isolate as *C. albicans* was supported by positive results from mass spectrometric identification with a score of 2.19 (Bruker Biotyper, Tricore Reference Laboratories), fungal ribosomal DNA ITS2 sequencing (Charles River, Accugenix FunITS), growth on chromagar, and filamentation assays. Similar assays were also completed for TRL031, confirming its identification as *C. glabrata*.

As a result of the marked increase in overall glucan exposure of both isolates, we hypothesized that the N-mannan in both samples must be deficient in specific features and would result in nanoscale alterations of glucan exposure sites similar to the mannosylation mutant strains described above. dSTORM imaging of TRL035 confirmed that the density of both single probe binding events and multi-exposures significantly increased as well as a significant increase in the median size of the exposures (Fig. 3A-F; SuppFig. 3A-E). The increase in both density and size of exposures mirrors what was observed in the *C. albicans* mannosylation mutants, specifically in the Ca*mnn2∆* mutant, leading us to hypothesize that TRL035’s N-mannan is deficient in some side chain features along with the acid labile fraction.

**Figure 3.**
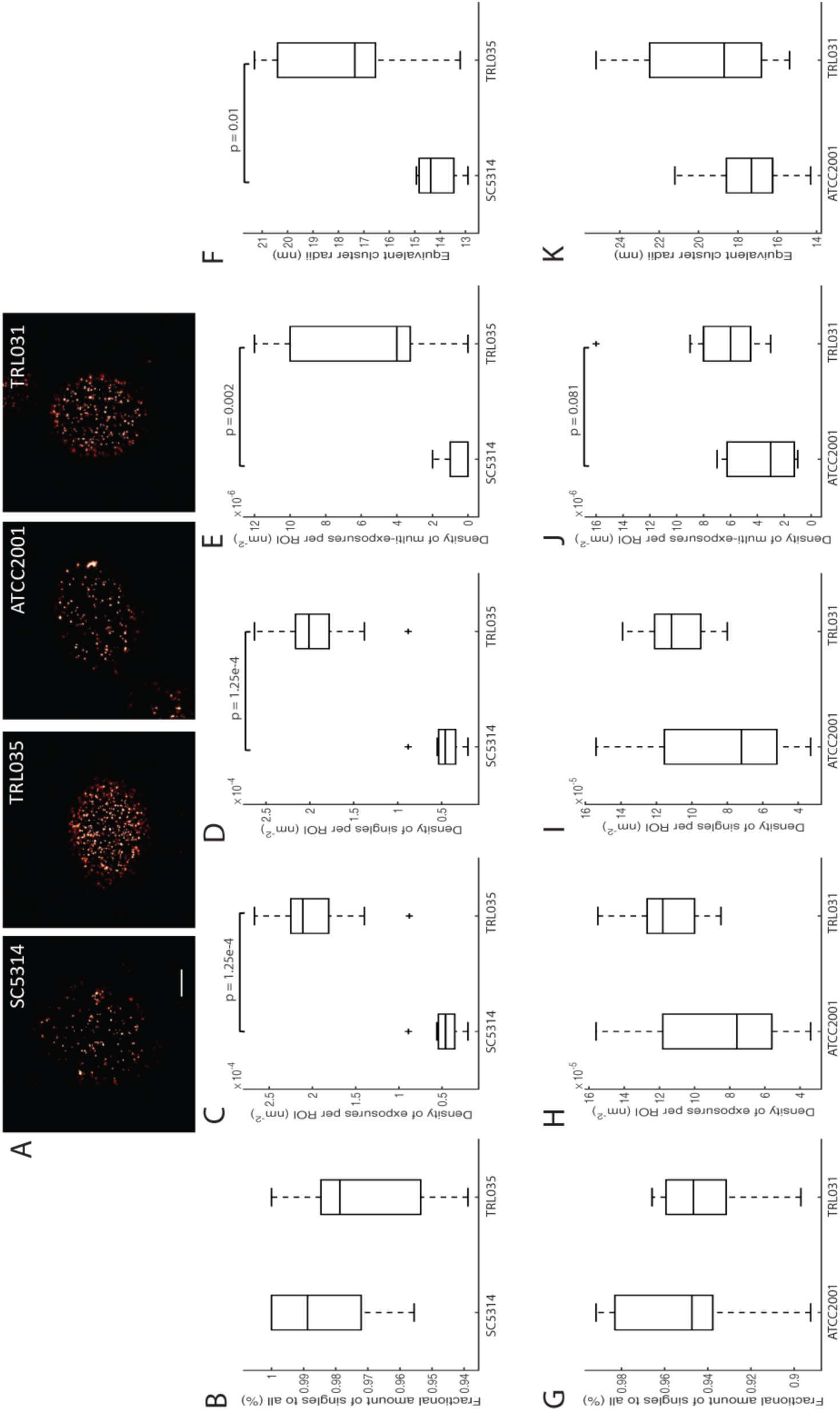
Clinical isolates nanoscopic β-glucan exposure characteristics are dissimilar from type strains. A) Representative dSTORM images of type strains and clinical isolates. Scale bar = 750 nm. B-F) Hierarchical clustering quantification of nanoscopic glucan exposure geometries on C. albicans strains. G-K) Hierarchical clustering quantification of nanoscopic glucan exposure geometries on C. glabrata strains. Box plots represent population of events found in ROI. Statistical significance was determined by Mann-Whitney U test with n ≥ 6 for each strain.

dSTORM imaging revealed that glucan multi-exposure density was slightly higher in TRL031 relative to ATCC2001 (Fig. 3J; SuppFig. 3I). No other nanoscale glucan exposure features were regarded to be significantly different (Fig. 3A,G-K SuppFig. 3F-J). When compared, the nanoscale exposure characteristics of TRL031 closely resembles the Cg*mnn2∆* mutant, leading to the hypothesis that TRL031s’ N-mannan is deficient in side chains and acid labile features.

### Physicochemical and structural analysis of N-mannan from *Candida* clinical isolates

To test if TRL035 and TRL031 have similar N-mannan features as the type strains, gel permeation chromatography with multiangle laser light scattering detection (GPC/MALLS) and ^1^H NMR were performed to determine the structure of the mannans. Mannan extraction was done on both isolates using a modified method from Li *et al.* 2009 (Li et al., 2009). TRL031’s mannoprotein molecular weight was found to be very similar to ATCC2001’s (Fig. 1J,4A). Poor yield of TRL035 mannoprotein, most likely due to small size, necessitated an alternate mild alkali extraction method. TRL035’s mannoproteins are found to be ~3-fold smaller than previously reported values for SC5314 (Fig. 1J,4A). Pronase treatment of TRL035 mannoprotein led to lower molecular weight mannan (66% decrease in molecular weight) and a narrower polydispersity of mannans (Fig.4A). In summary, GPC analysis shows that clinical isolate strain TRL035 produces significantly smaller N-mannans than type strain and is completely lacking high molecular weight mannoproteins characteristically found in *C. albicans*, whereas TRL031 produces similarly sized mannoproteins as characteristically found in *C. glabrata*.

We considered that the reduction in mannan size in TRL035 was likely because of lower structural complexity of the mannan, to confirm this hypothesis ^1^H NMR was used for analysis of TRL035 mannan structure. TRL035 has major resonances in the carbohydrate spectral region from 3.3 to 5.6 ppm, some of which provide information on mannan side chain structure. TRL035 mannan almost completely lacks the acid labile mannan resonances found abundantly in SC5314 mannan at 5.568, 5.183, 4.902, 4.855, 4.853, and 4.832 ppm (Fig. 4B,C). Loss of the acid labile mannan, containing a phosphodiester linkage, is anticipated to decrease the negative charge of the cell wall surface, which results in reduced binding of the polycationic dye Alcian Blue. Confirmation of TRL035 acid labile mannan deficiency was observed by the reduced Alcian Blue staining for TRL035 vs. SC5314 (Fig 4D). Because TRL035 has a clear deficiency the acid labile fraction, existing side chain resonances are assigned to the acid stable mannan fraction. TRL035 mannan’s signal at 5.068 ppm indicates, as expected, the presence of substituted α-(1,6)-linked backbone residues. Furthermore, the lack of a signal at 5.117 ppm for unsubstituted α-(1,6)-mannose suggests that the TRL035 mannan does not possess interspersed regions of acid stable side chain-free backbone (e.g., as was observed in Ca*mnn2*Δ, Ca*mnn2*Δ*/mnn26*Δ and Ca*mnn2* family sextuple mutant strains). The acid stable side chains are shown to be terminated in both α- and β-mannose units by the presence of signals at 5.280 ppm, 5.244 ppm and 5.002 ppm (Fig. 4B,C). Based on the NMR resonances found above, we constructed the most likely N-mannan structure for TLR035 (Fig. 4E).

**Figure 4.**
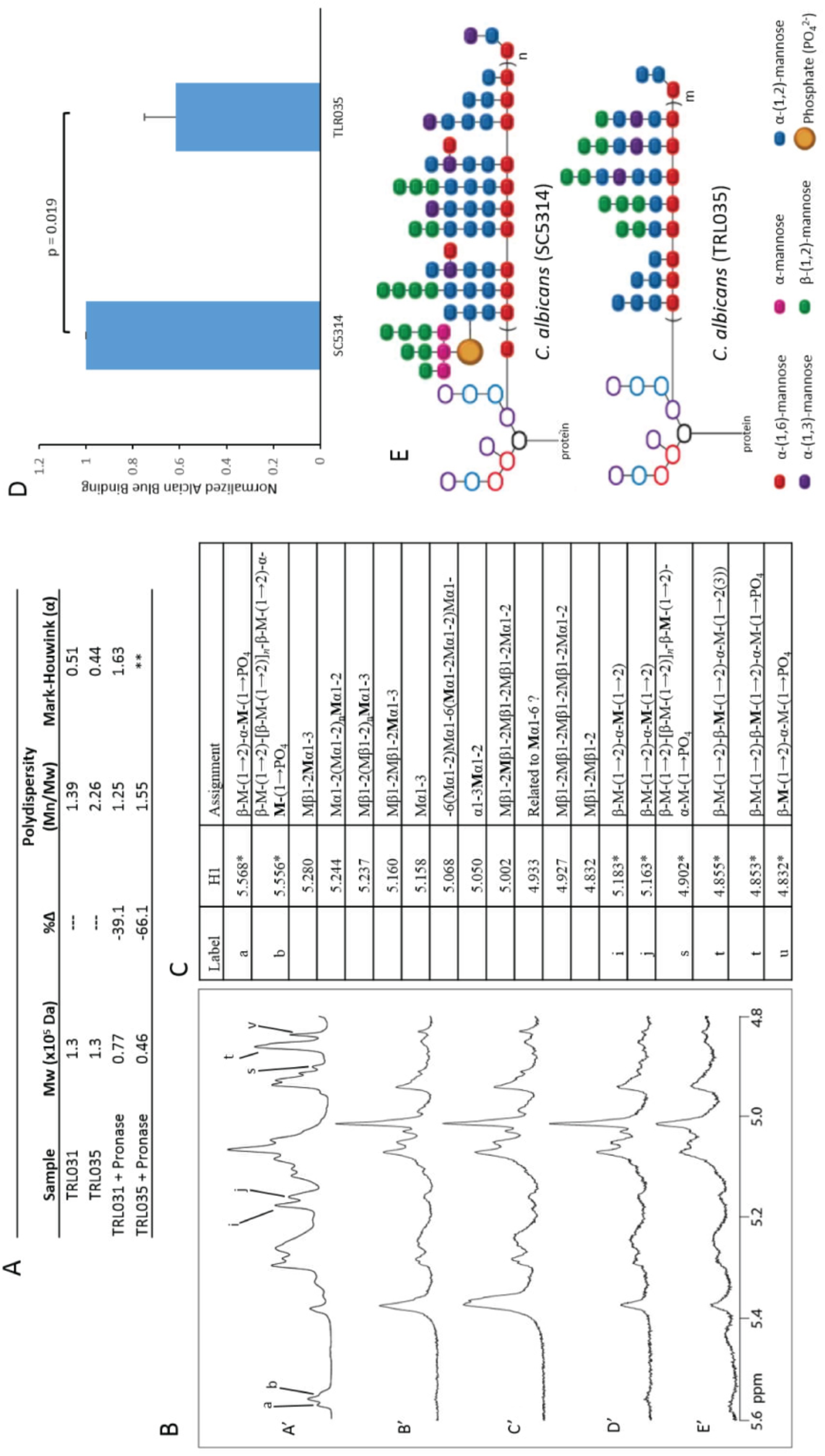
Physicochemical analysis shows loss of acid labile fraction in TRL035 N-mannan. A) GPC analysis of TRL031 and TRL035. ** denotes samples from which data could not be obtained due to low sample yield. B,C) NMR analysis of TRL035. B) Stacked spectral plot comparing SC5314 (A'), TRL035 (B',D'), and TRL035 pronase treatment (C',E'). B', C' and D', E' are from different cell wall isolations. C) Table of structural motif assignments for mannan. 1H chemical shifts with an asterisk are taken from Kruppa et al. 2011 (ref 39) to show where acid labile resonances are present in A'. D) Comparison of the amount of Alcian Blue bound per yeast. Statistical significance was determined by a single-tailed t-test with an n = 3. Values and error bars displayed are presented as normalized means and standard deviations. E) Proposed N-mannan structure of TRL035 based on NMR analysis compared to SC5314. Subscripts n and m correspond to an unspecified number of structural repeats > 1, and n > m.

Similar to TRL035, we also performed ^1^H NMR on TRL031. Compared to ATCC2001, TRL031 mannan lacks the resonances for the acid labile fraction of the mannan found in ATCC2001 at 5.555, 5.754, and 4.855 ppm (Fig. 5A,B). The Alcian Blue assay was attempted to confirm loss of acid labile fraction on TRL031 at the above chemical shifts (Fig. 5C). However, acid labile mannan is much less abundant in ATCC2001 than SC5314, making its loss in TRL031 a mild phenotype that was undetectable by the Alcian Blue assay. Acid stable resonances in ATCC2001 are also present in TRL031, showing that no additional loss of N-mannan features has occurred (Fig. 5D). TRL031 shows some similarities to the Cg*mnn2∆* mutant in that both strains have increased total glucan exposure along with increased size and density of the exposures that correlate with the loss of the acid labile fraction of the N-mannan.

**Figure 5.**
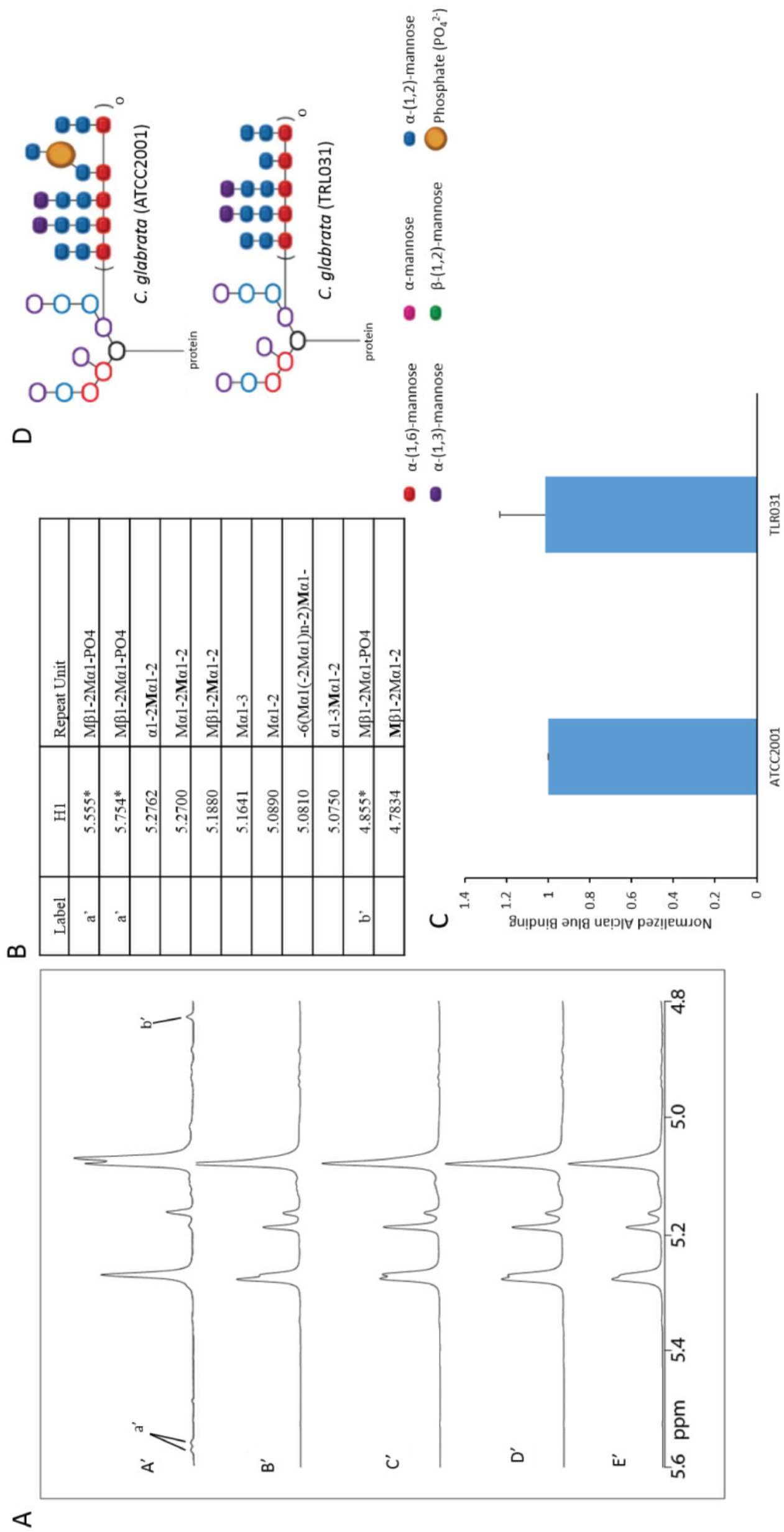
Physicochemical analysis shows loss of acid labile fraction in TRL031 N-mannan. A) Stacked spectral plots comparing ATCC2001 (A'), TRL031 (B',D'), and TRL031 pronase treatment (C',E'). B', C' and D', E' are from different cell wall isolations. C) Table of structural motif assignments for mannan. 1H chemical shifts with an asterisk are taken from West et al. 2011 (ref 9) to show where missing acid labile resonances are present in A'. C) Comparison of the amount of Alcian Blue bound per yeast. Statistical significance was determined by a single-tailed t-test with an n = 3. Values and error bars displayed are presented as normalized means and standard deviations. D) Proposed N-mannan structure of TRL031 based on NMR analysis compared to ATCC2001. Subscript o denotes an unspecified number of structural repeats > 1.

Taken together, the GPC and NMR data reveal an N-mannan structure in TRL035 that bears some similarities to *C. albicans* mannosylation mutant strains. More specifically, the loss of the acid labile fraction, which provides not only bulk but also negative charge, in *C. albicans* mannan is a feature that correlates with increased total glucan exposure as well as the size and density of nanoscale glucan exposure sites. The existence of clinical isolate strains that have acquired a mannan structure with high glucan exposure further indicates that regulation of glucan exposure through manipulation of mannan structure is not limited to artificial mutations, but can also be a part of the disease process in patients with candidiasis.

## Discussion

The outer layer of the yeast cell wall plays an essential role in protection and immune evasion. It is thought that the N-mannans of the yeast cell wall play a pivotal role in hindering Dectin-1 from binding to β-glucan. In this report, we define how specific N-mannan structural features affect immune evasion through the alteration of glucan exposure geometries at the molecular level. Our findings strongly implicate N-mannan side chain abundance and complexity in *C. albicans* and backbone length in *C. glabrata* as features that are important for masking glucan. In *C. albicans*, we found that as little as a single deletion in the MNN2 gene family can increase glucan exposure by over 4-fold (Fig. 1K, 2B). This could result from the loss of the negatively charged acid labile mannan, leading to less efficient glucan masking. Increasing the number of MNN2 gene family deletions increased the degree to which nanoscale glucan exposure phenotype changed. Among the *C. glabrata* mutants, the most significant changes to glucan exposure size and density arose from the mutants that have shortened backbones (Cg*mnn11∆* and Cg*anp1∆*). Consistent with this, Cg*mnn2∆*, which has the lowest Mw N-mannans of the *C. glabrata* mutant series, and only an unsubstituted mannan backbone, was able to provide glucan masking better than Cg*mnn11∆* and Cg*anp1∆,* which do retain some acid stable side chain structure.

Mannan masking of glucan in the cell wall can play a significant role in recognition of *C. albicans* and *C. glabrata* by the immune system. We showed that the ATCC2001 type strain of *C. glabrata* exhibits higher glucan exposure and more efficient phagocytic uptake by human monocyte-derived DCs than *C. albicans* (SC5314). Deletion of MNN2 family members is associated with increased glucan exposure in both C. albicans and C. glabrata. It is notable that previous studies have shown that these C. albicans MNN2 family mutants are hypovirulent in infection models, but the C. glabrata MNN2 mutants are hypervirulent (West et al., 2013)(Hall et al., 2013)(Rouabhia et al., 2005). Why does glucan exposure decrease virulence in *C. albicans* but increase virulence in *C. glabrata*? Hypovirulence of MNN2 family mutants in *C. albicans* may be explained with reference to enhanced host defense due to greater stimulation of glucan receptors such as Dectin-1 and CR3. On the other hand, *C. glabrata* has been shown to be able to survive and replicate within phagosomes (Seider et al., 2011), so more efficient entry into the phagosomal environment may promote survival, dissemination and evasion of immune detection for this pathogen.

Extending our work to patient samples, we found two clinical isolates with unusually high glucan exposures, TRL035 and TRL031. It is not yet clear how TRL035 acquired its high glucan exposure phenotype. It is possible that perturbation of one or more MNN2 family genes play a role, but other gene deletions cause increased glucan exposure. *cho1*Δ, *cek1*Δ and *kre5*Δ strains exhibit increased glucan exposure and Dectin-1 binding (Wheeler and Fink, 2006)(Davis et al., 2014)(Galan-Diez et al., 2010)(Herrero et al., 2004). *goa1*Δ mutants have recently been shown to have reductions in both β-(1,2)-linked mannose units and acid labile mannan (She et al., 2016), which bears some similarity to the TRL035 mannan phenotype. Future work will pursue molecular mechanistic explanations of mannan biosynthesis, glucan exposure and the pathological states associated with this and other high-glucan clinical *Candida* isolate strains.

The cause of high glucan exposure in TRL031 is also unclear. None of the *C. glabrata* mutants we investigated are structurally similar to TRL031. Based on GPC and NMR analysis, TRL031 does not seem to have a shortening of its α-(1,6)-mannose backbone nor a loss of side chains. The only difference detected in the N-mannan structure is the loss of the acid labile fraction, suggesting that the negatively charged fraction of the N-mannan plays a role in glucan masking. More research on mannosylation mutants is required to fully understand the effects of N-mannan biosynthesis on glucan exposure.

We consistently saw correlation of loss of acid labile mannan structure with increases in total glucan exposure and both the size and density of nanoscale glucan exposure features. It is therefore tempting to speculate that acid labile mannan is at least one important regulatory of glucan exposure nanostructure. Deletion of the CaMNN4 gene is associated with a more specific loss of acid labile mannan structure. Past reports have varied with respect to the impact of this defect on innate immune recognition of *C. albicans*. Hobson, et al examined a Ca*mnn4Δ* strain and reported no defect in virulence or phagocytic uptake at high yeast:phagocyte ratios (Hobson et al., 2004). However, McKenzie, et al did find that phagocytosis of a Ca*mnn4Δ* strain was significantly different from yeast with wild type mannan (McKenzie et al., 2010). Furthermore, Ueno, et al found that purified mannan from a Ca*mnn4Δ* strain induced significantly lower IL-6 and IL-12p40 responses *in vitro* relative to wild type mannan (Ueno et al., 2013). Likely impacts of reduction in acid labile mannan include altered adhesion in addition to impacts on glucan exposure, which may be reflected in the above studies. Our work suggests that future investigations into the role of CaMNN4 on glucan exposure on the micro- and nanoscale are indicated to further clarify the role of acid labile mannan in glucan masking.

Understanding the regulation of nanoscale glucan features by mannan structure is necessary for a more complete mechanistic appreciation of early events in pathogen recognition by innate immune cells. Dectin-1 signaling via its cytoplasmic hemITAM motif is thought to require ligand-dependent multimerization of the receptor (Inoue and Shinohara, 2014). The majority of glucan exposure sites on *C. albicans* cell walls are single receptor/ligand interaction sites (Lin et al., 2016) that may not support Dectin-1 multimerization. However, we have shown that exposure sites large enough for multimerization exist, which are comprised of clusters of multiple exposed glucan binding sites. We have also shown that perturbed mannan structures result in an increase in the surface density and individual size of these exposures. It is likely that these more complex exposed glucan nanostructures play critical roles in determining the multimerization of Dectin-1 and initiation of innate immune response to *Candida* species.

## Materials and Methods

### Fungal Culture

<u>*C. albicans.*</u> C. albicans (SC5314, ATCC, #MYA-2876), TRL035, and mannosylation mutants Ca*mnn2∆*, Ca*mnn2∆/mnn26∆*, and sextuple deletion mutants were grown in YPD broth in an orbital incubator at 30°C for 16 hours to reach exponential growth phase. All strains were then centrifuged and washed in PBS. Ca*mnn2∆*, Ca*mnn2∆/mnn26∆*, and sextuple deletion mutants were a gift from Dr. Neil Gow(Hall et al., 2013).

<u>*C. glabrata.*</u> C. glabrata (ATCC, #2001) and TRL031 were grown in YPD and mannosylation mutants Cg*mnn2∆,* Cg*mnn11∆,* and Cg*anp1∆* were grown in SD His^−^ media in an orbital incubator at 30°C for 16 hours to reach exponential growth phase. All strains were then centrifuged and washed in PBS. Cg*mnn2∆,* Cg*mnn11∆,* and Cg*anp1∆* were a gift from Dr. Ken Haynes(West et al., 2013).

### Clinical Strain Identification

TRL031 and TRL035 were grown on YPD agar plates for 48 hours at 30°C. Single colonies were transferred to Charles River (Newark, DE) on YPD slants. Both TRL031 and TRL035 were identified as *C. glabrata* and *C. albicans* respectively by FunITS at Charles River.

### Flow Cytometry

All strains of yeasts were treated similarly. After the PBS wash described above, yeast were counted on a hemocytometer (C-Chip Bulldog Bio, DHC-N01) to attain 3.5x10^6^ yeast total for primary and secondary staining. All strains were stained with 30µg/ml of Fc-hDectin-1a (Invivogen, fc-hdec1a) or 10µg/ml of murine anti-β-glucan IgM (clone BfDIV, gift of Biothera Pharmaceuticals) for 30 min at 25°C. The IgM probe was used for flow cytometric glucan exposure screening in clinical isolates. The Fc-hDectin-1a probe was used for all other glucan exposure experiments, including dSTORM. Yeasts were then centrifuged and washed with PBS three times. The secondary stain used to identify the primary probes was either AlexaFluor 488 conjugated anti-human IgG antibody (Invitrogen, A11013) for hDectin-1a or AlexaFluor 647 conjugated anti-murine IgM antibody (Invitrogen, A21240) for anti-β-glucan IgM. The secondary stain was added to the primary stained yeasts at 4µg/ml for 30 minutes at 25°C in the dark and then washed three times with PBS. Data were acquired using a BD LSRFortessa flow cytometer (BD Biosciences). All data presented was pooled from triplicate biological replicates of the samples.

### dSTORM Sample Preparation

All yeasts strains prior to probe application were fixed with 4% PFA at room temperature for 15 minutes followed by extensive washing. Yeasts were then counted in a similar manner as samples prepared for flow cytometry. AlexaFluor 647 (Invitrogen, A20006) was used to label the soluble Fc-hDectin-1a. AlexaFluor 647-NHS-ester and Fc-hDectin-1a were mixed at 10:1 molar ratio in PBS at pH 8 for 1 hour at 25°C in the dark. To purify the labeled proteins, Zeba Desalting Columns (Thermo Scientific, 7-kDA MWCO, 89882) were used. Degree of labeling was found to be 1:1 molar ratio determined by absorption spectroscopy. Yeasts were then stained with the AlexaFluor 647 conjugated Fc-hDectin-1a at 30µg/ml for 30 min at 25°C in the dark and then washed three times in PBS. Prior to imaging, the fixed/labeled samples were embedded in 1% agar on the coverslip to minimize motion during image acquisition.

### dSTORM imaging

Imaging methods were duplicated from Lin et al. 2016. In summary, the agar embedded samples had an imaging buffer applied to them for 15 minutes prior to imaging. Imaging buffer contained 50mM Tris, 10% (wt/vol) glucose, 10 mM NaCl, 40µg/ml catalase (Sigma-Aldrich, C40), 500µg/ml glucose oxidase (Sigma-Aldrich, G0543), and 10 mM cysteamine (Sigma-Aldrich, M9768), pH 8. Data acquisition was on an Olympus IX-71 microscope equipped with an objective-based TIRF illuminator using an oil-immersion objective (PlanApo N, 150×/1.45 NA; Olympus) in an oblique illumination configuration. Sample excitation was done with a 637nm laser (Thorlabs, laser diode HL63133DG), with custom-built collimation optics. To ensure that minimal drift occurred during data acquisition, a self-registration algorithm was implemented.

### Super-resolution data fitting

To fit probe positions from raw dSTORM data, we used a maximum likelihood-based algorithm from Smith et al. 2010(Smith et al., 2010). Our positional fitting parameters included: normal fitting with a Cramer-Rao lower bound (CRLB) based fit precision threshold of 0.2 pixels or 22 nm. This suppressed the contribution of positionally inaccurate fits arising from rare events where multiple dyes were emitting simultaneously in a diffraction-limited volume. Fluorophores identified in time frames with a frame gap of 1 to four frames were combined into a single, improved fit when the spatial proximity passed a hypothesis test with 0.05 level of significance. Frame connection suppressed overrepresentation of fits from dyes with unusually long on-times, which is expected from the exponential distribution of dye on-times.

### Super-resolution cluster analysis

Analysis of nanostructure in our dSTORM datasets proceeded in three steps: estimating the positions of single emitter probes, identifying spatial clusters of exposed glucan labeled by probes, and quantifying feature characteristics thus identified.

<u>Hierarchical Single Emitter Hypothesis Test (H-SET).</u> Single probes can be observed more than once, with positional error, in dSTORM datasets. This contributes some overestimation of probe density and feature size. The purpose of the H-SET algorithm is to estimate the positions of single emitter probes and minimize such errors in subsequent cluster identification steps.

All image data was run through pass 1 of H-SET, a top-down hierarchical clustering algorithm implemented in MATLAB that collapses clusters of observations of blinking fluorophores into single estimates of the true locations (localizations) of the fluorophores(Lin et al., 2016). Essentially all collapses occur in pass 1, which requires only one user specified parameter, a level of significance for the hypothesis test (LoS = 0.01 was used), to make its decisions.

Pass 2 of H-SET determines clustering of exposures by using some known clustering algorithm, which for distance or density based methods (e.g., hierarchical or DBSCAN(Ester et al., 1996)) typically depends on two parameters: a distance metric, epsilon, the maximal distance between two points within a cluster, and the minimal number of points (minPts) required to form a cluster. These parameters define the type of cluster that will be detected, and it is often impossible to know these values *a priori* for biological structures. We optimized the choice of epsilon and minPts in a data driven fashion.

At any given parameterization of epsilon and minPts, spurious clusters can be detected in a spatially random distribution of points once the density of points becomes high enough. We sought to parameterize epsilon and minPts such that detection of spurious clustering in simulated random distributions would be minimized, while still remaining sensitive enough to detect clusters in datasets. Therefore, we optimized epsilon and minPts using the mean density of localizations in our densest datasets, which was 649 localizations per ROI for the *C. albicans* sextuple mutant dataset (eight ROIs over five images).

To determine values for epsilon and minPts, H-SET pass 2 hierarchical clustering (MATLAB linkage function) was performed on the sextuple mutant dataset for a range of epsilons and minPts (20-40 nm by 1 nm and 5-15, respectively), computing the number of multiglucan exposure sites, N_multi_, for each combination averaged over all the ROIs. These values were compared with the mean results from n = 25 random simulations generated over the same ranges of epsilon and minPts, the simulated localizations being distributed uniformly and randomly over the ROI, and the number of which were computed from a Poisson distribution with mean based on the average number of localizations determined from the sextuple mutant data. The biggest difference between N_multi_ for the two datasets, remaining sensitive to detection of multiglucan exposure sites for the experimental data but minimizing their spurious detection for the random simulated data, produced unique values for epsilon and minPts: 20 nm and 6, respectively, for *C. albicans*, and 26 nm and 5, respectively, for *C. glabrata*. Supplemental Figure 4 shows 3D plots of N_multi_ as a function of epsilon and minPts for the two datasets, (SuppFig. 4A) experimental and (SuppFig. 4B) random simulated, as well as their difference (SuppFig. 4C).

<u>Voronoï tessellation-based segmentation.</u> To verify that the trends observed were independent of the clustering algorithm employed, the analyses were recomputed using the Voronoï-based algorithm of Levet et al. (Levet et al., 2015), using our own MATLAB implementation. The algorithm proceeds by constructing a Voronoï diagram of image points (SuppFig. 5A, localizations). A Voronoï diagram is a partitioning of a region based on a set of seeds (points) within that region such that any location in a given cell is closer to the seed used to generate that cell than the seeds used to form the other cells. See Supplemental Figure 5A. Polygon 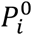 corresponds to seed (point) *s*_i_ and has area 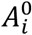. First rank neighbors are those polygons 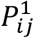 sharing edges with 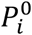, which when combined together with 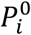 form the rank 1 polygon 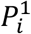. The rank 1 area 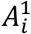 is then the sum of the areas 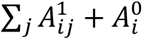. From these, densities δ^*k*^ (k is the rank) can be computed by dividing the number of points in 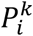 by 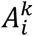. For further clarification, see Levet et al., 2015, Figure 1(Levet et al., 2015).

At this point, object segmentation (clustering) is initiated. For example, if δ is the average density of the data distribution, then clustering proceeds by merging all rank k polygons that satisfy

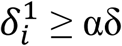

where α > 0 (k = 1 and α = 2 are typical choices in Levet et al.'s analyses).

The polygons satisfying the above criterion that are adjacent to each other form potential clusters of the seeds that generated them, and are merged into single clusters if the final collections have the user specified minimum number of points, minPts. Supplemental Figure 5B displays the clustering results based on the partitioning shown in Supplemental Figure 5A, which was based on one ROI of the sextuple mutant dataset.

Similar to what was done previously, pass 2 H-SET was performed using the Voronoï based algorithm with k = 1 over ranges of α (1-3 by increments of 0.25) and minPts (3-15), computing once again the number of multiglucan exposure sites. These were compared, as before, with random simulated results, choosing values for α and minPts where the difference between the experimental data and the random simulated results was the greatest at an α and minPts: 2 and 5 respectively for *C. albicans* and 2 and 4, respectively, for *C. glabrata*.

<u>Computer code.</u> Software used for data analysis and simulation is accessible at <u>http://stmc.health.unm.edu/tools-and-data/index.html</u>, Image Analysis Tools, MATLAB Clustering Classes.

### Alcian Blue binding analysis

Alcian Blue binding assay was adapted from Hobson et al. 2004(Hobson et al., 2004). Yeasts were grown for 16 hours, washed, counted to achieve a number of 1x10^7^ cells/ml and suspended in 25µg/ml of Alcian Blue (Sigma Aldrich, A5268) in 0.02 M HCl for 25 minutes at room temperature before being centrifuged and pelleted. Supernatant was then measured in a plate reader at A_600_. Alcian Blue concentrations were calculated against a standard curve. Alcian Blue binding (*x* A_600_ units/cell) was found by calculating the initial A_600_ (*u*), the final A_600_ (*v*), and (*n*) the number of cells stained. These values were input into the formula: *x* = (*u – v) / n*. Data is presented as normalized to the mean of type strain Alcian Blue binding. All data presented was pooled from triplicate biological replicates of the samples.

### GPC analysis

<u>Cultivation of fungal strains.</u> *C. albicans* strains were grown overnight (16hr) in 2L of YPD (2% Dextrose, 2% Peptone, 1% Yeast Extract). The cells were harvested by centrifugation at 5000 x g and stored at −20°C until they were needed for mannan extraction.

<u>Isolation of cell wall mannan.</u> Mannan was prepared from cells using a modified extraction, which we have previously utilized in Li et al., 2009 (Li et al., 2009). Briefly, cells were delipidated with acetone, and the pellet disrupted by bead beating in distilled H_2_O. The extracted cells were then autoclaved for 3 hr, and the extract centrifuged. The supernatant was split in half for each sample with one-half treated with 100 mg pronase (heated to 70°C for 30 min to kill glycosidic activity) for 18 hr at 37°C. The supernatant was subjected to Fehling precipitation. The resulting precipitate from the pronase treated and untreated sample fractions were then dissolved in 3 M HCl and the carbohydrate precipitated and copper removed with an 8:1 methanol:acetic acid solution. The solution was washed several times until the remaining precipitate was white to colorless. The precipitate was dried, resuspended in dH_2_O, and the pH adjusted to 6.5-7.0 after which the solution was lyophilized for storage. The yeast strains mannans were provided by Dr. Mike Kruppa as a lyophilized powder.

<u>Isolation of cell wall mannan by mild alkali bath.</u> Mannans were isolated using a new method (Kwofie *et al*, manuscript in preparation) that utilized a mild alkali extraction with boiling for 20 min. The extracted samples were harvested by centrifugation and neutralized. The neutralized material then was split in two and one-half treated with 50 mg/sample pronase (heat treated 25 min at 65°C to remove glycosidic activity) for 18 hours at 37°C in order to reduce the protein component of the mannoprotein. All samples were subjected to carbohydrate precipitation by addition of four volumes of methanol and samples were allowed to sit while the material settled. Once the material had settled, the supernatant was decanted and the remaining methanol blown off by blowing air over the sample. The dried sample was then resuspended in dH_2_O, frozen, lyophilized and stored at −20°C until it was analyzed.

<u>Gel permeation chromatographic analysis of *C. albicans* cell wall mannan/mannoprotein</u>. The molecular weight, polydispersity, polymer distribution and Mark-Houwink (α) values were derived from gel permeation chromatography (GPC) with a Viscotek/Malvern GPC system consisting of a GPCMax autoinjector fitted to a TDA 305 detector (Viscotek/Malvern, Houston, TX). The TDA contains a refractive index detector, a low angle laser light scattering detector, a right angle laser light scattering detector, an intrinsic viscosity detector and a UV detector (λ = 254 nm). Three Waters Ultrahydrogel columns, i.e, 1200, 500 and 120, were fitted in series (Waters Corp. Milford, MA). The columns and detectors were maintained at 40°C within the TDA 305. The system was calibrated using Malvern pullulan and dextran standards. The mobile phase was 50mM sodium nitrite. The mobile phase was filter sterilized (0.45 µm into a 5 L mobile phase reservoir). Mannan samples were dissolved (3 mg/ml) in mobile phase (50 mM sodium nitrite, pH 7.3). The samples were incubated for ~60 min at 60°C, followed by sterile filtration (0.2 µm) and injected into the GPC (200 µL). Sample recovery was routinely >98%. The data were analyzed using Viscotek OmniSec software v. 4.7.0.406. dn/dc was calculated using the OmniSec software (v. 4.6.1.354). dn/dc for the mannan samples was determined to be 0.145. The data were analyzed using a single peak assignment in order to obtain an average Mw for the entire polymer distribution. Replicate analysis of calibration standards indicated reproducibility of + or - 3%, which is well within the limits of the technique.

### NMR analysis

NMR data acquisition and analysis is based on methods from Lowman et al. 2011(Lowman et al., 2011). In summary, ^1^H NMR spectra for mannan were collected on a Bruker Avance III 600 NMR spectrometer using a DCH cryoprobe operating at 345 °K (72 °C) in 5mm NMR tubes as reported previously(Lowman et al., 2011). Mannan (about 10 mg) was dissolved in about 600 µL DH_2_O (Cambridge Isotope Laboratories, 99.8+% deuterated). Chemical shift referencing was accomplished relative to TMSP at 0.0 ppm. NMR spectra were collected and processed as follows: 256 scans, 2 pre-scans, 32,768 data points, 20 ppm sweep width centered at 6.2 ppm, and 1 s pulse delay. Spectra were processed using exponential apodization with 0.3 Hz line broadening and the JEOL DELTA software package (version 5.0.4.4) on a MacBook Pro (operating system OSX Yosemite 10.10.5). COSY spectra were collected as 2048 by 256, processed as 1024 by 1024, 8 pre-scans, 16 scans, sweep width 9 ppm centered at 4.7 ppm, and relaxation delay 1.49 sec. Processing was accomplished with sine apodization in both dimensions using TopSpin (version 3.5 pl 3) on the MacBook Pro.

### Statistics

Bar graphs were used to display the median of the integrated intensity of β-glucan on the yeast cells as measured by flow cytometry. Error bars represent the standard deviation of the data. Student's t-test, single-tail was used to determine statistical significance. Box plots were used to display population distributions for dSTORM data sets. These plots depict a median (solid line) within a box representing the interquartile range (25th to 75th percentile of the population). The whiskers below and above the box represent the 1st and 99th percentiles of the data set. Crosses are used to depict data points that fell outside of the whiskers. Mann-Whitney U test was used to determine statistical significance of the box plots and was implemented by MATLAB functions.

## Acknowledgements

We appreciate and acknowledge Janet Oliver for helpful scientific discussion and review of the manuscript as well as the members of the Neumann laboratory. We thank Dr. Karissa Culbreat and Christen Griego-Fullbright for their assistance with providing clinical isolate strains. This research was supported by the University of New Mexico Center for Spatiotemporal Modeling of Cell Signaling (STMC; NIH P50GM085273, AKN and KAL) and R01AI116894 (AKN). MSG was supported by fellowships from STMC and NIH T32 training grant (NIH T32 AI1007538) during the course of this work. This work was also supported, in part, by Quillen College of Medicine, East Tennessee State University, through NIH GMR0153522 (DLW), GMR01119197 (DLW), and GMR01083016 (DLW), NIH R15AI109581 (MDK), and C06RR030651.

## Author Contributions

Conceived and designed experiments: MSG and AKN. Performed the experiments: MSG, MDK, DLW, DWL, and JMY. Data analysis: MSG, AKN, MJW, KAL, DLW, DWL, and HCP. Contributed reagents/materials/analysis tools: MJW. Manuscript production: MSG, AKN, and MJW.

## Disclosure

The authors disclose that there are no financial interests or conflicts of interest.

## Supplemental Figure Legends

Supplemental Figure S1. Nanoscopic β-glucan exposure characteristics are determined by N-mannan structure. A-E) Voronoï tessellation-based segmentation quantification of nanoscopic glucan exposure geometries on C. albicans strains. F-J) Voronoï tessellation-based segmentation quantification of nanoscopic glucan exposure geometries on C. glabrata strains. Box plots represent population of events found in ROI. Statistical significance was determined by Mann-Whitney U test with n ≥ 6 for each strain.

Supplemental Figure S2: Clinical isolates expose more β-glucan compared to type strains. A) Comparison of glucan exposure of SC5314 to different medical isolates. Only TRL035 had a significant increase in glucan exposure in comparison to SC5314. Data was pooled from triplicate biological repeats with an n ≥ 10000 events per sample. B) Representative images of type strains and medical isolates with varying amounts of β-glucan exposed. Scale bar = 10 µm. C,D) Comparison of the integrated intensity of β-glucan. Statistical significance was determined by a single-tailed t-test with n ≥ 20000 events per strain. Values and error bars displayed in C and D are presented as normalized medians and standard deviations.

Supplemental Figure S3. Clinical isolates nanoscopic β-glucan exposure characteristics are dissimilar from type strains. A-E) Voronoï tessellation-based segmentation quantification of nanoscopic glucan exposure geometries on C. albicans strains. F-J) Voronoï tessellation-based segmentation quantification of nanoscopic glucan exposure geometries on C. glabrata strains. Box plots represent population of events found in ROI. Statistical significance was determined by Mann-Whitney U test with n ≥ 6 for each strain.

Supplemental Figure S4. Number of multiglucan exposure sites, Nmulti, as a function of the clustering parameters epsilon and minPts. A) Densest experimental data (sextuple mutant dataset) averaged over 8 ROIs from 5 images. B) Random data averaged over n = 25 simulations. C) The difference between A and B. Epsilon is denoted as "E".

Supplemental Figure S5. Voronoï based clustering of one ROI of the sextuple mutant dataset for rank k = 1 and α = 2. A) Voronoï tessellation of the dataset, where the colors of the polygons indicate their relative rank k density with respect to the overall density of the ROI. Displayed points are the seeds used to form the tessellation, where red indicates seeds that are considered potentially clustered as the corresponding polygonal region has rank k density ≥ αδ, where δ is the average density within the entire ROI. B) Clusters produced from (A) during pass 2 of H-SET, indicated by green boundary lines. The lines were constructed by the MATLAB boundary command using a shrink factor of 0.5 (the default value). The cyan dots are the original observations, while the dark blue dots are the collapsed localizations produced by H-SET. Red boundary lines, when present, group clusters of localizations that were collapsed in pass 2 of H-SET.

